# Adolescent Binge Ethanol Exposure Confers Lasting Alcohol Tolerance across a Cumulative Ethanol Challenge in Adulthood: Involvement of Proinflammatory HMGB1 Signaling

**DOI:** 10.1101/2025.04.17.649339

**Authors:** Fulton T. Crews, Ryan P. Vetreno

## Abstract

**Background:** Epidemiological studies suggest heavy adolescent binge drinking is strongly associated with later development of an alcohol use disorder (AUD). Alcohol tolerance (i.e., an acquired reduction in acute alcohol responsivity) is a universally recognized symptom of AUD, but the direct contribution of adolescent binge drinking to adult alcohol tolerance is poorly understood.

**Methods and Materials:** To investigate the contributions of adolescent binge ethanol exposure to lasting acquisition of acute tolerance, we used our ethanol response battery (ERB) to assess intoxication rating, hypothermia, motor coordination, and balance across cumulative ethanol doses (i.e., 0.0, 0.5, 1.0, 2.0, and 3.0 g/kg) in adult female Wistar rats following adolescent intermittent ethanol (AIE), lipopolysaccharide (LPS), and glycyrrhizic acid treatment following AIE.

**Results:** We report AIE, which models human adolescent binge drinking, confers lasting alcohol tolerance across cumulative ethanol doses and blunts ethanol-induced increases in proinflammatory HMGB1 plasma levels. Adolescent LPS (1.0 mg/kg, i.p.) treatment, which mimics AIE-induced HMGB1-mediated neuroinflammation, induces adult alcohol tolerance and blunts HMGB1 release across cumulative ethanol doses on the ERB. Assessment of proinflammatory HMGB1 involvement in AIE-induced acquisition of lasting alcohol tolerance revealed that post-AIE administration of the HMGB1 inhibitor glycyrrhizic acid reversed the AIE-induced acquisition of alcohol tolerance in adulthood.

**Conclusions:** These data reveal that (1) adolescent binge drinking confers long-lasting low ethanol responsivity, (2) proinflammatory neuroimmune activation contributes to the development of alcohol tolerance, and (3) blockade of proinflammatory HMGB1 signaling reverses AIE-induced acquisition of alcohol tolerance in adulthood. These findings suggest a potential mechanistic target for the development of novel therapeutics for the treatment of AUD.

## Introduction

Alcohol tolerance is a commonly endorsed symptom of alcohol use disorder (AUD) (1, 2) that likely contributes to increased binge drinking and risk of alcohol dependence. A low level of alcohol responsivity (i.e., tolerance) predicts later development of binge drinking and increased risk for lifelong alcohol problems (3–5). Epidemiological studies report that an adolescent age of drinking onset is a significant risk factor for later development of an AUD (6–11). Adolescents have low alcohol responsivity and binge drinking is common during adolescence. Binge drinking, which involves rapid consumption of multiple drinks (4-5 drinks over a 2-hr period), increases the rewarding effects of alcohol contributing to the development of AUD (12–17). Preclinical binge ethanol studies find rapid ethanol-induced dopamine release, a known proxy for reward, in the nucleus accumbens; this release declines as ethanol plateaus, with repeated doses leading to a new peak in dopamine (18), consistent with binge drinking to keep blood ethanol rising to intoxication. Preclinical (19) and clinical studies (20) find adolescent drinking impacts brain development, which increases risks for AUD. In this study, we systematically investigated the impact of adolescent intermittent ethanol (AIE), a preclinical rodent model of human adolescent binge drinking, on adult behavioral and physiological responsivity across multiple measures during an acute cumulative ethanol challenge (21).

Assessment of alcohol responsivity is affected by metabolism, time after ethanol, and rising or falling blood ethanol levels (BECs). One consistent finding is that, compared to adults, adolescents have a low response to alcohol (22). Using the tilting plane, White and colleagues (23, 24) reported that acute ethanol elicited low responsivity in adolescent rats (P30-P65) relative to adults (P70-P105), with adolescents developing chronic tolerance, but not adults with identical ethanol exposure. Similarly, chronic intermittent ethanol treatment of male rats during adolescence altered loss of righting reflex (LORR) sleep time, but not the aerial righting reflex, in adulthood (25). We recently compared adolescent (postnatal [P]40) and adult (P85) male and female rat ethanol responsivity using an ethanol response battery (ERB) cumulative dose response that assesses intoxication, hypothermia, rotarod motor coordination, tilting plane balance, and LORR. Across cumulative ethanol doses and endpoints, adolescent males and females were significantly less sensitive to ethanol than adults (22). Although the mechanisms underlying this age-associated differential response to alcohol are unknown, the ERB provides a full ethanol dose response curve, allowing determination of whether AIE confers alcohol tolerance in adulthood.

Prior AIE studies found increased adult alcohol drinking and preference, increased anxiety and pain sensitivity, and cognitive deficits, all of which could increase risk for development of AUD (18). Studies of adult acute ethanol induction of immediate early genes following AIE found reduced cFos+IR in prefrontal cortex across multiple doses. consistent with tolerance (26). However, tolerance to alcohol is poorly understood and linked to multiple neurobiological systems (1, 21, 27). Chronic ethanol exposure studies support excitatory and inhibitory synaptic changes as contributing to development of physical dependence, and the hyper-excitability of alcohol withdrawal (28) has been proposed to contribute to alcohol-induced tolerance. Although our AIE model does not induce physical dependence, it does alter adult behaviors that increase risks for AUD development (19). Previous studies find AIE induces long-lasting changes in brain gene expression through epigenetic mechanisms that persist long after the last ethanol exposure (29, 30). We also find AIE increases adult expression of multiple proinflammatory cytokines including HMGB1, which is a nuclear cytokine known to be released by acute ethanol and activate epigenetic changes in neurons (31–33), proinflammatory activation of glia (34, 35), and persistently alter behavior (36–39). Emerging studies suggest ethanol-induced proinflammatory signaling alters synaptic excitability (40) and contributes to increased alcohol drinking and preference (41–44), consistent with proinflammatory signaling altering synapses that could change alcohol responsivity and contribute to alcohol tolerance. While accumulating evidence suggests a role for alcohol-induced neuroinflammation in alterations in alcohol drinking and AUD risk, neuroinflammatory involvement in alcohol tolerance is unknown. To investigate the contributions of adolescent binge ethanol and potential neuroimmune mechanisms of AIE-induced developmental acquisition of alcohol tolerance in adulthood, we used our ethanol response battery (ERB) to assess adult acute ethanol responsivity across cumulative ethanol doses following AIE (Experiment 1), an adolescent neuroinflammatory lipopolysaccharide (LPS) challenge (Experiment 2), and post-AIE treatment with the HMGB1-specific inhibitor glycyrrhizic acid (Experiment 3). Since our prior ERB studies did not find major sex differences, we focused on females in the current investigation. The results from these studies support the hypothesis that proinflammatory HMGB1 signaling induces adult ethanol tolerance that is reversible, consistent with proinflammatory activation as a key target to reduce risk for AUD (45).

## Methods and Materials

### Animals

Female Wistar rats bred at the University of North Carolina at Chapel Hill were used in this study. Subjects were housed in same-treatment pairs in a temperature-(20°C) and humidity-controlled vivarium on a 12 hour/12 hour light/dark cycle (light onset at 7:00 AM) and provided *ad libitum* access to food and water. This study was conducted in an AAALAC-accredited facility in strict accordance with NIH regulations for the care and use of animals in research. All experimental procedures reported in this study were approved by the Institutional Animal Care and Use Committee at UNC.

### Adolescent Intermittent Ethanol (AIE) Paradigm

On P21, female Wistar rats (N=56) were randomly assigned to either (i) AIE (n=28 [Experiment 1: n=8; Experiment 3: n=20) or (ii) water control (CON; n=28 [Experiment 1: n=8; Experiment 3: n=20) treatments. To minimize the impact of litter variables, no more than one subject from a given litter was assigned to a single experimental condition. From P25 to P54, AIE subjects received a single daily intragastric (i.g.) administration of ethanol (EtOH; 5.0 g/kg, 25% EtOH, w/v) on a 2-day on/2-day off schedule, and CON subjects received an equivalent volume of water on an identical schedule as previously described (46). Subjects in Experiment 1 underwent ERB assessment on P75 and subjects in Experiment 3 were assessed on the ERB on P95 (**Figure 1A**).

**Figure 1.**
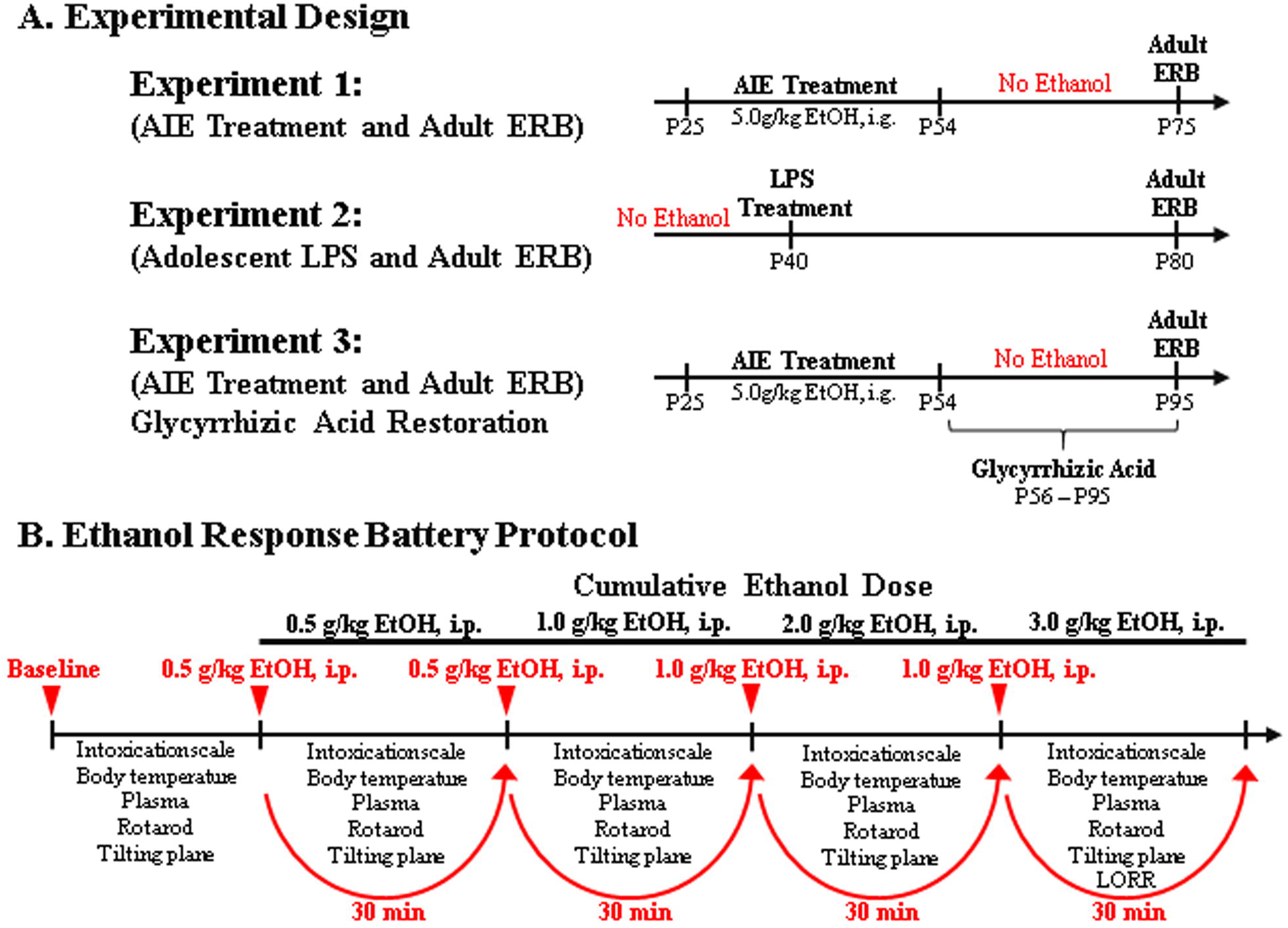
Schematic of the experimental design and ethanol response battery (ERB) protocol. **(A)** In Experiment 1, female Wistar rats received either control (CON) or adolescent intermittent ethanol (AIE) treatment from P25–P54 followed by ERB assessment on P75. In Experiment 2, ethanol-naïve rats received a single dose of lipopolysaccharide (LPS, 1.0 mg.kg, i.p.) on P40 and were assessed on the ERB on P80. In Experiment 3, rats received CON or AIE treatment from P25–P54 followed by glycyrrhizic acid (HMGB1 inhibitor) treatment from P56–P95, and assessment of ERB performance on P95. Body weights were assessed throughout experimentation. Tail blood was collected from AIE- and CON-treated subjects to assess blood ethanol concentrations (BECs) using a GM7 Analyzer (Analox; London, UK) 1 hr after treatment on P38 (AIE/VEH: 141±22 mg/dL; AIE/Glyz Acid: 170±18 mg/dL) and P54 (AIE/VEH: 187±24 mg/dL; AIE/Glyz Acid: 195±40 mg/dL). **(B)** Each trial of the ERB consisted of (1) the 6-point behavioral intoxication rating scale, (2) body temperature assessment, (3) tail blood collection, (4) accelerating rotarod assessment (Experiments 1 and 2 only), (5) tilting plane assessment, and (6) loss of righting reflex (LORR) following the final ethanol dose. The ERB was conducted during a cumulative ethanol dose-response challenge. Following baseline ERB assessment, rats received ethanol doses (0.5, 0.5, 1.0, 1.0 g/kg, i.p.) approximately 30 min apart for cumulative ethanol doses of 0.5, 1.0, 2.0, and 3.0 g/kg with the ERB initiated 15 min following each ethanol dose.

### Adolescent Lipopolysaccharide (LPS) Treatment

In Experiment 2, female Wistar rats (N=16; n=8/group) received either a single injection of LPS (1.0 mg/kg, i.p. in sterile 0.9% saline; E. Coli, serotype 0111:B4; Sigma-Aldrich, Cat. #L3024) or a comparable volume of vehicle on P40. Subjects underwent ERB assessment on P80 (**Figure 1A**).

### Glycyrrhizic Acid Treatment

Adult CON- and AIE-treated rats in Experiment 3 received post-AIE treatment with glycyrrhizic acid (1.0 g/L, P.O.; Sigma-Aldrich; Cat. #50531) or vehicle (water) in cage water bottles from P56 to P95. Subjects in the glycyrrhizic acid condition consumed an average of 37.4 mL/day with an approximate average consumption of 140 mg/kg glycyrrhizic acid per day.

### Ethanol Response Battery (ERB)

The ERB was performed as described previously (47, 48) and methodology is detailed in **Supplement 1**. Animals underwent ERB assessment beginning at 8:00 AM during the light cycle consisting of a baseline assessment followed by four consecutive cumulative doses of ethanol to assess ethanol responsivity across a broad range of BECs. The four consecutive doses of ethanol (i.e., 0.5, 0.5, 1.0, 1.0 g/kg, i.p.) produce cumulative ethanol doses of approximately 0.5, 1.0, 2.0, and 3.0 g/kg (see **Figure 1B**). Each subsequent ERB assessment was conducted approximately 30 min apart, each initiated 15 min after ethanol dosing. The ERB consisted of (1) the 6-point behavioral intoxication rating scale, (2) body temperature assessment, (3) tail blood collection, (4) rotarod motor coordination (Experiments 1 and 2 only), (5) tilting plane balance, and (6) LORR following the final ethanol dose.

### HMGB1 ELISA

Blood samples collected during the ERB were centrifugated (2,000×g for 20 min) and plasma assessed for HMGB1. HMGB1 ELISA was performed according to the manufacturer’s protocol (IBL International, Hamburg, Germany, Cat. #ST51011).

### Immunohistochemistry, Microscopic Quantification, and Image Analysis

Following the conclusion of ERB assessment in Experiment 3, brain tissue was collected for assessment of HMGB1 and pNF-κB p65 expression in the motor cortex as previously reported (49–51). More details on immunohistochemistry, antibodies, and microscopy methodology are described in **Supplement 1**.

### Statistical Analysis

Statistical analysis was performed using SPSS (IBM; Chicago, IL) and GraphPad Prism 8 (San Diego, CA). BECs, intoxication rating, body temperature, rotarod, tilting plane, and HMGB1 plasma data were first assessed using the Shapiro-Wilk test to determine normality of data distribution. Data with a normal distribution were assessed using parametric repeated measure ANOVAs with post-hoc Bonferroni corrections or Šidák’s MCT as described in the Results. Data that was not normally distributed was assessed using the non-parametric Mann-Whitney test. Ordinal data (i.e., intoxication rating) was assessed using the Mann-Whitney test (Experiments 1 and 2) or the Kruskal-Wallis H test (Experiment 3). Immunohistochemistry data (Experiment 3) was assessed using a 2 × 2 ANOVA with post-hoc Tukey’s HSD performed as necessary. Baseline threshold assessments were analyzed using ANOVA with post-hoc Bonferroni corrections. Pearson Chi-Square was used to analyze LORR data. All values except intoxication rating are reported as mean ±SEM. The intoxication rating data is reported as median with the interquartile range.

## Results

### Adolescent binge ethanol exposure confers long-lasting low ethanol responsivity across a cumulative ethanol challenge in adulthood

Assessment of BECs during cumulative ethanol doses revealed remarkably similar progressive increases, ranging from approximately 72 mg/dL (0.5 g/kg) to 307 mg/dL (3.0 g/kg), that did not differ as a function of AIE treatment (**Figure 2A** and **Figure S1A in Supplement 1**). Across cumulative ethanol doses, both CON- and AIE-treated adult (i.e., P75) rats evidenced increasing intoxication scores with rising BECs (**Figure 2A/B**). Prior AIE treatment significantly decreased intoxication scores at 1.0 g/kg (*p*=0.027, Mann-Whitney), 2.0 g/kg (*p*=0.0002, Mann-Whitney), and 3.0 g/kg (*p*=0.027, Mann-Whitney) relative to age-matched CONs (**Figure 2B**). Across cumulative ethanol doses, both CON- and AIE-treated adult rats evidenced increasing hypothermia (main effect of Dose: *p*<0.0001; **Figure 2C** and **Figure S1B in Supplement 1**). Post-hoc analyses revealed AIE blunted the hypothermic response at 0.5 g/kg (*p*=0.011, Bonferroni), 1.0 g/kg (*p*=0.007, Bonferroni), 2.0 g/kg (*p*=0.002, Bonferroni), and 3.0 g/kg (*p*=0.001, Bonferroni) relative to age-matched CONs (**Figure 2C**). Further, adult CONs evidenced a significant baseline hypothermic threshold shift at the 1.0 g/kg dose (*p*=0.0009, Bonferroni) whereas AIE-treated animals evidenced a significant baseline threshold difference at the 3.0 g/kg dose (*p*=0.0004, Bonferroni). At the 3.0 g/kg dose, core body temperatures decreased by 2.7°C in CONs compared to a 1.2°C decrease in AIE-treated animals. Time on the accelerating rotarod, which provides an index of motor coordination, progressively decreased as ethanol dose increased (**Figure 2D** and **Figure S1C in Supplement 1**). Prior AIE treatment significantly increased time on the rotarod at 2.0 g/kg (*p*=0.021, Mann-Whitney) relative to CONs. Adult CONs evidenced a significant baseline rotarod threshold shift at 1.0 g/kg (*p*=0.037, Bonferroni) whereas AIE-treated animals evidenced a significant baseline threshold shift at the 2.0 g/kg dose (*p*<0.0001, Bonferroni). Angle of slide on the tilting plane, which provides a measure of balance, dose-dependently decreased across cumulative ethanol doses in adult CON- and AIE-treated rats (main effect of Dose: *p*<0.0001; **Figure 2E** and **Figure S1D in Supplement 1**). Post-hoc analyses revealed that AIE treatment increased the angle of slide at 1.0 g/kg (*p*=0.0005, Bonferroni), 2.0 g/kg (*p*=0.0004, Bonferroni), and 3.0 g/kg (*p*=0.0002, Bonferroni) relative to CONs. Further, adult CONs evidenced a significant baseline tilting plane threshold shift at the 1.0 g/kg dose (*p*=0.043, Bonferroni) whereas AIE-treated animals evidenced a significant baseline threshold shift at the 3.0 g/kg dose (*p*<0.0001, Bonferroni). Assessment of LORR revealed that all adult CON subjects (n=8) displayed LORR whereas only 4 of the 8 adult AIE-treated rats demonstrated a LORR (*p*=0.021, Pearson Chi-Square; **Figure 2F**). Thus, adult AIE-treated rats evidenced lower intoxication scores, decreased hypothermia, and diminished impairment of motor coordination and balance across a broad range of acute cumulative ethanol doses.

**Figure 2.**
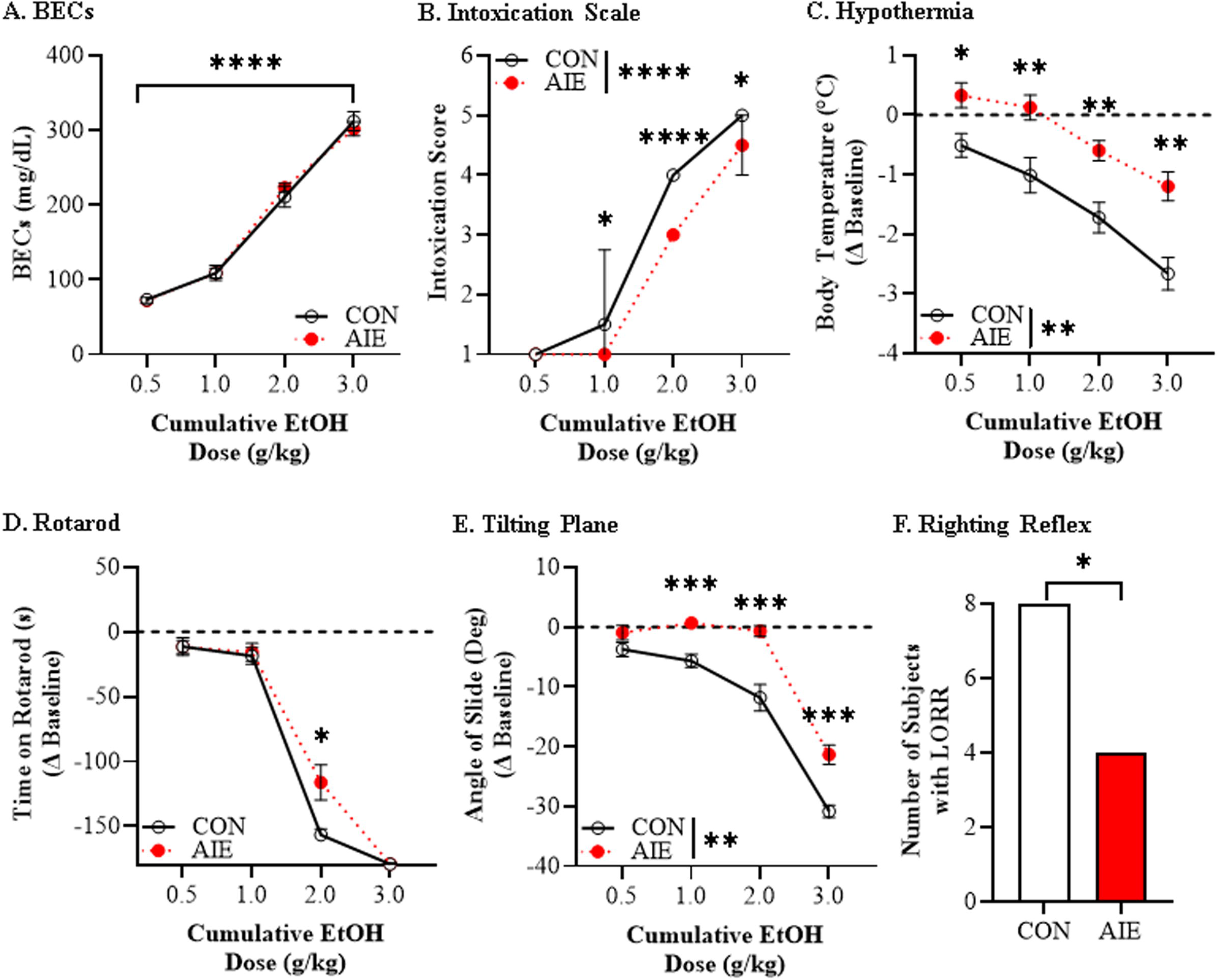
Adolescent intermittent ethanol (AIE) confers long-lasting low ethanol responsivity on the ethanol response battery (ERB) in adulthood. **(A)** Assessment of BECs 15 min after each ethanol dose across the cumulative ethanol challenge during the ERB revealed a dose-dependent increase in BECs (main effect of Dose: *F*[3, 42]=473.5, *p*<0.0001) that did not differ as a function of AIE treatment. **(B)** Assessment of intoxication rating across the cumulative ethanol challenge revealed increasing intoxication scores across treatment conditions. Prior AIE treatment significantly decreased adult intoxication scores at 1.0 g/kg (*U*=16.0, *p*=0.027, Mann-Whitney test), 2.0 g/kg (*U*=0.00, *p*=0.0002, Mann-Whitney test), and 3.0 g/kg (*U*=16.0, *p*=0.027, Mann-Whitney test) relative to age-matched CONs. **(C)** Across adult CON- and AIE-treated female rats, assessment of body temperature during the cumulative ethanol challenge revealed increasing hypothermia with rising BECs (main effect of Dose: *F*[3, 42]=105.2, *p*<0.0001). Further, AIE blunted the hypothermic response at 0.5 g/kg (Bonferroni correction: *p*=0.011), 1.0 g/kg (Bonferroni correction: *p*=0.007), 2.0 g/kg (Bonferroni correction: *p*=0.002), and 3.0 g/kg (Bonferroni correction: *p*=0.001) relative to age-matched CONs. **(D)** Assessment of rotarod motor coordination revealed a dose-dependent impairment in time to remain on the rotarod across cumulative ethanol doses. Relative to age-matched adult CONs, adult AIE-treated rats spent significantly more time on the rotarod at the 2.0 g/kg dose (*U*=16, *p*=0.021, Mann-Whitney test). **(E)** Assessment of tilting plane balance in adult female rats revealed a dose-dependent reduction in the angle of slide across adult CON- and AIE-treated rats (main effect of Dose: *F*[3, 42]=191.2, *p*<0.0001). Prior AIE treatment increased the angle of slide at 1.0 g/kg (Bonferroni correction: *p*=0.0005), 2.0 g/kg (Bonferroni correction: *p*=0.0004), and 3.0 g/kg (Bonferroni correction: *p*=0.0002) relative to age-matched CONs. **(F)** Assessment of loss of righting reflex (LORR) following the final dose of ethanol (i.e., 3.0 g/kg) revealed all adult CON subjects displays LORR whereas 4 of the 8 adult AIE-treated rats demonstrated a LORR (*X^2^*(1, *N*=16)=5.3, *p*=0.021, Pearson Chi Square). n=8 subjects/condition. Data are presented as mean ±SEM or median with interquartile range. \**p*<0.05, \*\**p*<0.01, \*\*\**p*<0.001, \*\*\*\**p*<0.0001.

High mobility group box 1 (HMGB1) is a proinflammatory cytokine-like protein released by ethanol in neuronal, microglial, and astrocyte cultures as well as in forebrain and hippocampal slice culture models (49, 52–56). We next assessed plasma levels of HMGB1 in adult animals following AIE treatment during the cumulative ethanol challenge. Baseline HMGB1 plasma levels ranged from approximately 25 to over 100 ng/mL, with baseline increases in AIE-treated animals relative to CONs (*p*=0.004, Student’s *t* test; **Figure 3A**). Across the ERB, we observed a dose-dependent increase in plasma HMGB1 levels in CON- and AIE-treated rats (main effect of Dose: *p*=0.0003; **Figure 3B**). Post-hoc analyses revealed that AIE blunted plasma HMGB1 levels at 0.5 g/kg (*p*=0.089, Bonferroni), and significantly reduced plasma HMGB1 at 1.0 g/kg (*p*=0.001, Bonferroni), 2.0 g/kg (*p*=0.046, Bonferroni), and 3.0 g/kg (*p*=0.043, Bonferroni) relative to CONs. Interestingly, adult CONs evidenced a significant baseline plasma HMGB1 threshold shift at the 2.0 g/kg dose (*p*=0.016, Bonferroni) whereas baseline threshold shifts were not observed in the adult AIE-treated animals. These findings are consistent with AIE altering acute ethanol-induced release of HMGB1.

**Figure 3.**
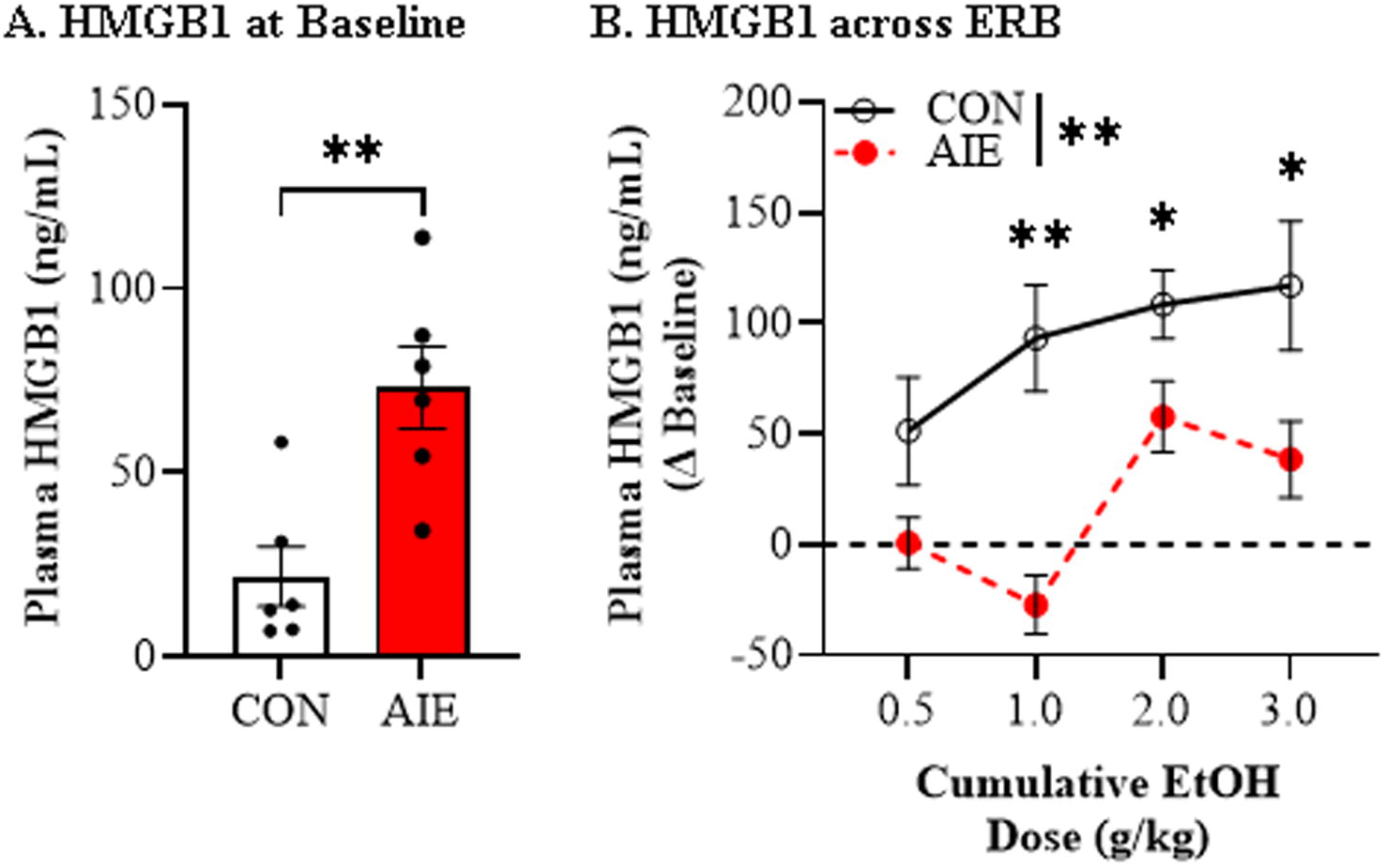
Adolescent intermittent ethanol (AIE) alters adult plasma high mobility group box 1 (HMGB1) responsivity to a cumulative ethanol challenge during ethanol response battery (ERB) assessment in adulthood. **(A)** Assessment of baseline plasma HMGB1 levels prior to ERB assessment in adulthood revealed a persistent AIE-induced 3.4-fold increase (Student’s *t* test: *t*(10)=3.7, *p*=0.004) relative to age-matched CONs. **(B)** Across cumulative ethanol doses of the ERB, plasma levels of HMGB1 increased in a dose-dependent manner in CON- and AIE-treated adult rats (main effect of Dose: *F*[3, 30]=8.7, *p*=0.0003). In adult rats with a history of AIE exposure, ERB assessment revealed a trend for reduced plasma HMGB1 at 0.5 g/kg (Bonferroni correction: *p*=0.089), and a significant reduction of plasma HMGB1 at 1.0 g/kg (Bonferroni correction: *p*=0.001), 2.0 g/kg (Bonferroni correction: *p*=0.046), and 3.0 g/kg (Bonferroni correction: *p*=0.043) relative to age-matched CONs. n=6 subjects/condition. Data are presented as mean ±SEM. \**p*<0.05, \*\**p*<0.01.

### An adolescent neuroimmune challenge with LPS confers long-lasting low ethanol responsivity to a cumulative ethanol challenge in adulthood

We have previously shown that AIE induces proinflammatory gene expression, including HMGB1, throughout the adult brain (39, 57–59). Similar to AIE, LPS causes release of HMGB1 and induction of proinflammatory genes in brain (52, 60). To determine if adolescent proinflammatory gene induction alters adult acute alcohol responsivity, we treated adolescent rats with LPS on P40 and assessed ERB performance 40 days later on P80. Across the ERB, BECs progressively increased (main effect of Dose: *p*<0.0001) and did not differ as a function of adolescent LPS treatment (**Figure 4A**). Adolescent LPS administration significantly blunted adult intoxication scores at doses of 1.0 g/kg (*p*=0.033, Mann-Whitney) and 2.0 g/kg (*p*=0.015, Mann-Whitney) relative to CONs (**Figure 4B**). Assessment of body temperature during adult ERB testing revealed an increasing hypothermic response across cumulative ethanol doses (main effect of Dose: *p*<0.0001; **Figure 4C** and **Figure S2B in Supplement 1**), similar to AIE treatment. Post-hoc analyses revealed that adolescent LPS administration decreased the hypothermic response at 1.0 g/kg (*p*=0.024, Bonferroni), blunted the hypothermic response at 2.0 g/kg (*p*=0.057, Bonferroni), and decreased the hypothermic response at 3.0 g/kg (*p*=0.002, Bonferroni) relative to CONs. Further, adult CONs evidenced a significant baseline hypothermic threshold shift at the 0.5 g/kg dose (*p*=0.009, Bonferroni) whereas LPS-treated animals evidenced a significant baseline threshold shift at the 2.0 g/kg dose (*p*<0.0001, Bonferroni). At the 3.0 g/kg dose (∼300 mg/dL), core body temperatures decreased by 3.4°C in CONs compared to a 2.2°C decrease in AIE-treated animals. Interestingly, while adult CON- and LPS-treated animals evidenced dose-dependent decreases in time on the rotarod (**Figure 4D** and **Figure S4C in Supplement 1**), we did not observe alterations in rotarod motor coordination due to adolescent LPS exposure. Assessment of balance on the tilting plane revealed a dose-dependent reduction in angle of slide across cumulative ethanol doses in CON- and LPS-treated rats (main effect of Dose: *p*<0.0001; **Figure 4E** and **Figure S4D in Supplement 1**). Interestingly, adolescent LPS decreased overall tilting plane impairment in adulthood relative to CONs (main effect of Treatment: *p*=0.007). Further, adult CONs evidenced a significant baseline tilting plane threshold shift at the 2.0 g/kg dose (*p*=0.002, Bonferroni) whereas LPS-treated animals evidenced a significant baseline threshold shift at the 3.0 g/kg dose (*p*<0.0001, Bonferroni). Assessment of LORR revealed that all adult CON subjects displayed LORR whereas 6 of the 8 adult LPS-treated rats demonstrated a LORR (*p*=0.131, Pearson Chi-Square; **Figure 4F**).

**Figure 4.**
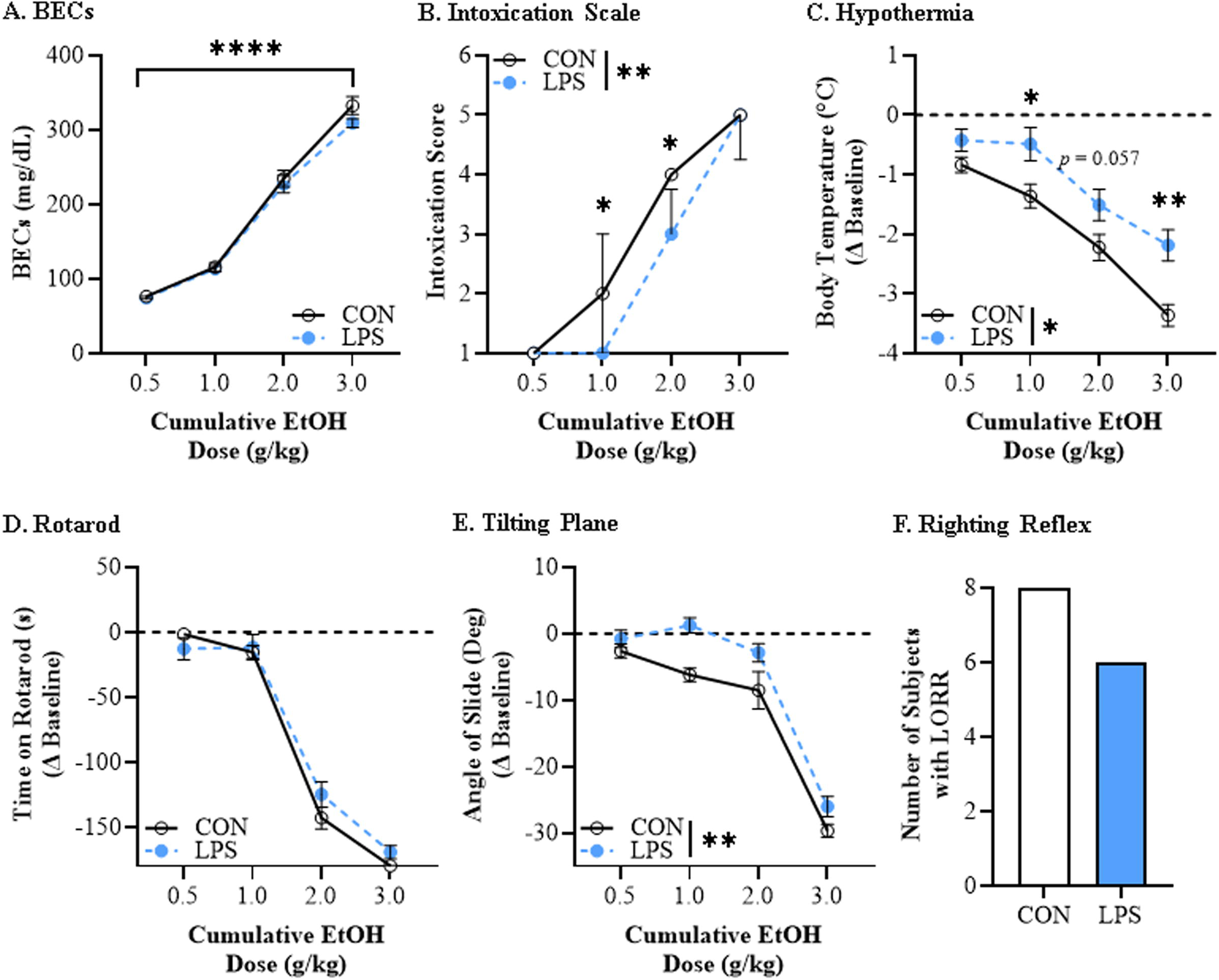
Adolescent lipopolysaccharide (LPS) exposure confers long-lasting low ethanol responsivity on the ethanol response battery (ERB) in adulthood. **(A)** Assessment of BECs 15 min after each ethanol dose across the cumulative ethanol challenge during the ERB revealed a dose-dependent increase in BECs (main effect of Dose: *F*[3, 42]=529.2, *p*<0.0001) that did not differ as a function of prior LPS exposure. **(B)** Intoxication rating assessment across the cumulative ethanol challenge revealed increasing intoxication scores in adult CON- and LPS-treated rats. Adolescent LPS treatment decreased adult intoxication scores at doses of 1.0 g/kg (*U*=14.5, *p*=0.033, Mann-Whitney test) and 2.0 g/kg (*U*=12, *p*=0.015, Mann-Whitney test) relative to age-matched CONs. **(C)** Adult CON- and LPS-treated rats evidenced an increasing hypothermic response across cumulative ethanol doses (main effect of Dose: *F*[3, 42]=143.0, *p*<0.0001). Prior adolescent LPS treatment decreased the hypothermic response at 1.0 g/kg (Bonferroni correction: *p*=0.024), blunted the hypothermic response at 2.0 g/kg (Bonferroni correction: *p*=0.057), and decreased the hypothermic response at 3.0 g/kg (Bonferroni correction: *p*=0.002) relative to age-matched CONs. **(D)** Assessment of motor coordination on the rotarod revealed that latency to fall decreased, regardless of treatment condition, across cumulative ethanol doses (main effect of Dose: *p*<0.0001) while prior LPS exposure did not affect rotarod performance. **(E)** Assessment of tilting plane balance in adult female rats revealed a dose-dependent reduction in the angle of slide across adult CON- and LPS-treated rats (main effect of Dose: *F*[3, 42]=189.1, *p*< 0.0001). Adolescent LPS treatment led to an overall decrease in tilting plane impairment in adulthood relative to age-matched CONs (main effect of Treatment: *F*[1, 14]=10.0, *p*=0.007). **(F)** Assessment of loss of righting reflex (LORR) following the final dose of ethanol (i.e., 3.0 g/kg) revealed all adult CON subjects displays LORR whereas 6 of the 8 adult LPS-treated rats demonstrated a LORR (*X^2^*(1, *N*=16)=2.3, *p*=0.131, Pearson Chi Square). n=8 subjects/condition. Data are presented as mean ±SEM or median with interquartile range. \**p*<0.05, \*\**p*<0.01, \*\*\*\**p*<0.0001.

Plasma HMGB1 levels at baseline did not differ between CON- and LPS-treated adult rats (**Figure 5A**). Across cumulative ethanol doses, we observed a dose-dependent increase in plasma HMGB1 levels (main effect of Dose: *p*=0.017; **Figure 5B**). Post-hoc analyses revealed an adolescent LPS-induced reduction in plasma HMGB1 levels at ethanol doses of 1.0 g/kg (*p*=0.042, Bonferroni), 2.0 g/kg (*p*=0.046, Bonferroni), and 3.0 g/kg (*p*=0.004, Bonferroni) relative to CONs. These findings, coupled with lower intoxication ratings, decreased hypothermia, and blunted motor impairments, suggest that an adolescent neuroimmune challenge, perhaps through a HMGB1-mediated neuroinflammatory mechanism, confers lasting alcohol tolerance similar to AIE treatment.

**Figure 5.**
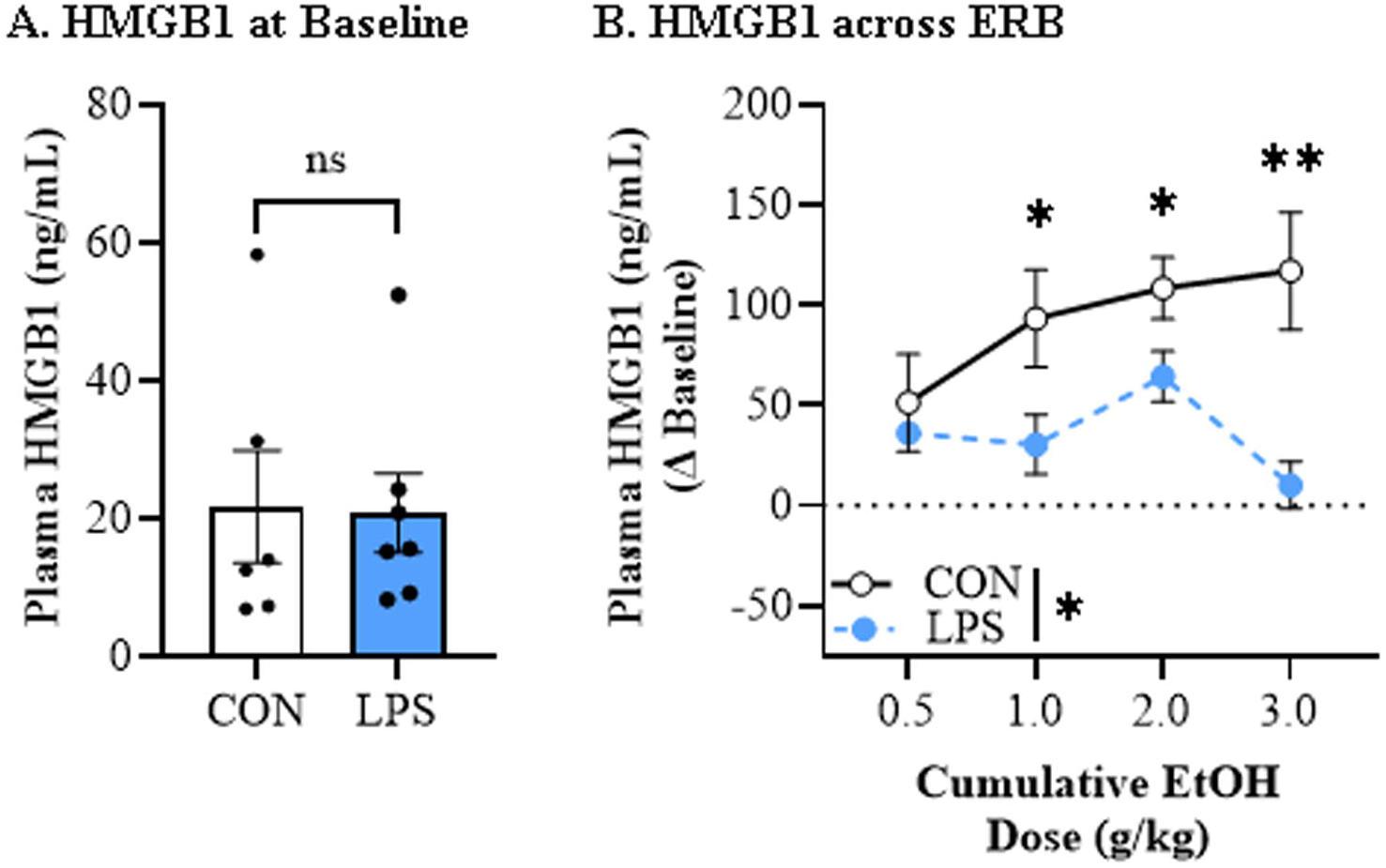
Adolescent lipopolysaccharide (LPS) exposure alters adult plasma high mobility group box 1 (HMGB1) responsivity to a cumulative ethanol challenge during ethanol response battery (ERB) assessment in adulthood. **(A)** Baseline HMGB1 plasma levels did not differ between treatment groups prior to ERB assessment in adulthood (Student’s *t* test: *p*=0.93). **(B)** Across cumulative ethanol doses of the ERB, plasma levels of HMGB1 increased in a dose-dependent manner in CON- and adolescent LPS-treated adult rats (main effect of Dose: *F*[3, 33]=3.9, *p*=0.017). In adult rats with a history of adolescent LPS exposure, ERB assessment revealed a reduction in plasma HMGB1 levels at ethanol doses of 1.0 g/kg (Bonferroni correction: *p*=0.042), 2.0 g/kg (Bonferroni correction: *p*=0.046), and 3.0 g/kg (Bonferroni correction: *p*=0.004) relative to age-matched CONs. n=6 subjects/condition. Data are presented as mean ±SEM. \**p*<0.05, \*\**p*<0.01.

### The HMGB1 inhibitor glycyrrhizic acid reverses AIE-induced long-lasting low ethanol responsivity to a cumulative ethanol challenge in adulthood

Previous studies suggest HMGB1 signaling contributes to AIE-induced proinflammatory gene induction that persists into adulthood (19, 61–63). Glycyrrhizic acid is an HMGB1 inhibitor that binds to a central pocket on HMGB1, antagonizing activation of multiple proinflammatory receptors (64). To determine if AIE-induced alterations in ERB tolerance involve proinflammatory HMGB1 signaling, we administered glycyrrhizic acid from P56 (48 hr post-AIE) until ERB assessment on P95 (**Figure S3A in Supplement 1**). Assessment of BECs revealed remarkably similar progressive increases (main effect of Dose: *p*<0.0001) that did not differ across groups (**Figure 6A**). Similar to Experiment 1, intoxication rating differed significantly across treatment groups with adult vehicle-treated AIE rats demonstrating lower intoxication scores at ethanol doses of 0.5 g/kg (*p*=0.003, Kruskal-Wallis), 1.0 g/kg (*p*=0.0002, Kruskal-Wallis), and 2.0 g/kg (*p*=0.0003, Kruskal-Wallis) relative to CONs. While glycyrrhizic acid did not affect intoxication ratings in CONs, it normalized the AIE-induced low intoxication rating to CON levels at ethanol doses of 0.5 g/kg (*p*=0.012, Kruskal-Wallis), 1.0 g/kg (*p*=0.017, Kruskal-Wallis), and 2.0 g/kg (*p*=0.055, Kruskal-Wallis) relative to vehicle-treated AIE animals (**Figure 6B**). Regardless of treatment condition, all rats evidenced increasing hypothermia (main effect of Dose: *p*<0.0001), whereas follow-up analyses revealed a blunted hypothermic response in vehicle-treated AIE animals at ethanol doses of 0.5 g/kg (*p*=0.018, Bonferroni) and 1.0 g/kg (*p*=0.003, Bonferroni) relative to CONs. Glycyrrhizic acid did not affect hypothermic responses in CONs, but reversed the AIE-induced blunting of hypothermia at ethanol doses of 0.5 g/kg (*p*=0.015, Bonferroni) and 1.0 g/kg (*p* = 0.017, Bonferroni) relative to vehicle-treated AIE animals (**Figure 6C** and **Figure S3B in Supplement 1**). Balance assessment on the tilting plane was similarly reduced across treatment groups (main effect of Dose: *p*<0.0001), whereas a history of AIE exposure increased the angle of slide at 0.5 g/kg (*p*<0.0001, Bonferroni), 1.0 g/kg (*p*<0.0001, Bonferroni), and 2.0 g/kg (*p* < 0.0001, Bonferroni) relative to vehicle-treated CONs. While glycyrrhizic acid did not affect tilting plane performance in CONs, it restored cumulative ethanol-induced impairments at doses of 0.5 g/kg (*p*=0.001, Bonferroni), 1.0 g/kg (*p*=0.009, Bonferroni), and 2.0 g/kg (*p*=0.005, Bonferroni) relative to vehicle-treated AIE animals (**Figure 6D** and **Figure S3C in Supplement 1**). In this cohort, assessment of LORR revealed no differences across treatment conditions (**Figure 6E**). Thus, similar to Experiment 1, adult vehicle-treated AIE rats evidenced increased adult tolerance to acute ethanol intoxication that was reversed by treatment with the HMGB1 inhibitor glycyrrhizic acid.

**Figure 6.**
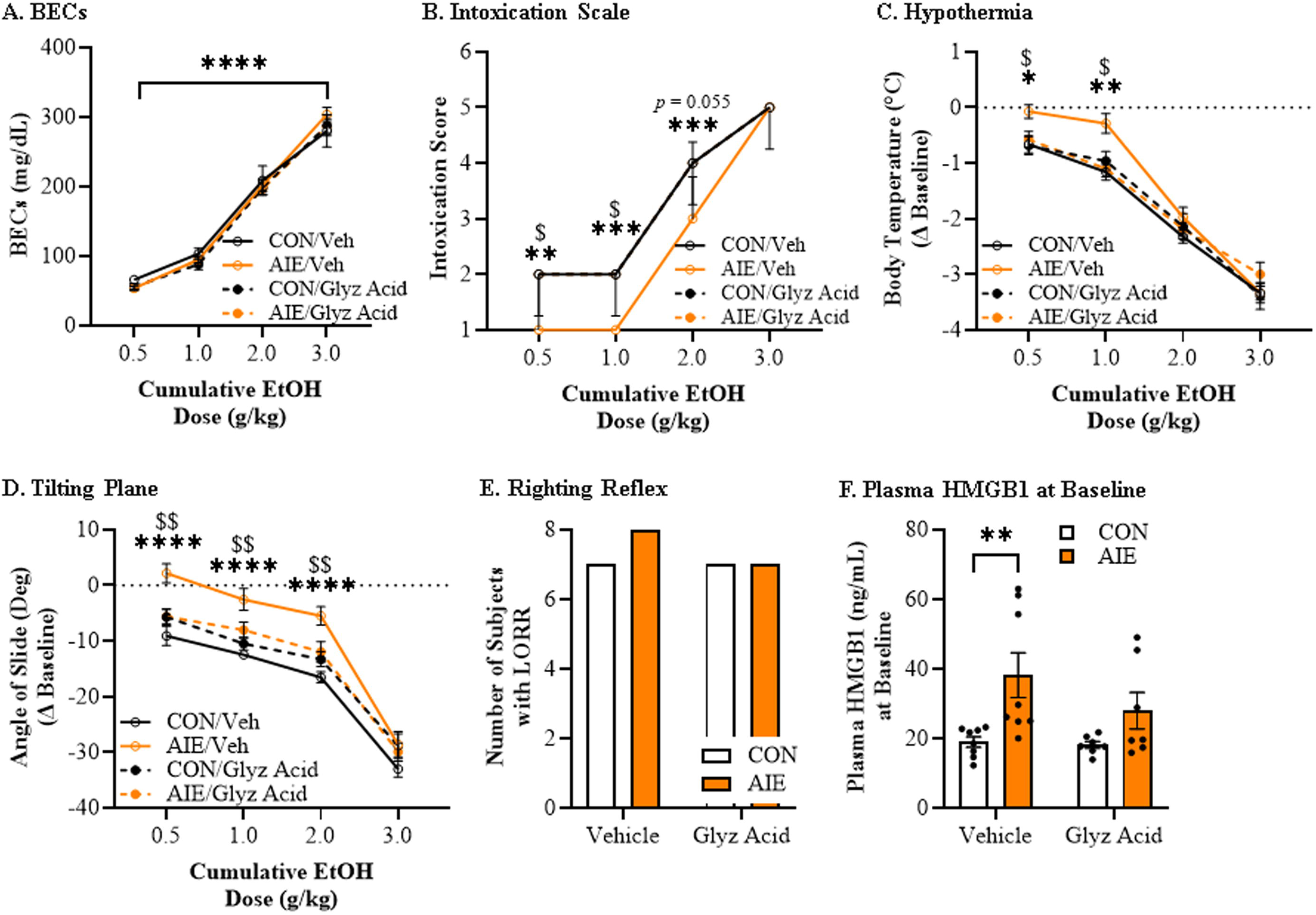
Adult treatment with the HMGB1 inhibitor glycyrrhizic acid reverses adolescent intermittent ethanol (AIE)-induced acquisition of long-lasting low ethanol responsivity to a cumulative ethanol challenge in adulthood. **(A)** Assessment of BECs 15 min after each ethanol dose across the cumulative ethanol challenge during the ERB revealed a dose-dependent increase in BECs (main effect of Dose: *F*[3, 84]=518.9, *p*<0.0001) that did not differ as a function of AIE or glycyrrhizic acid treatment. **(B)** Assessment of intoxication rating across the cumulative ethanol challenge revealed increasing intoxication scores, regardless of treatment condition. Prior AIE treatment significantly decreased adult intoxication scores at 0.5 g/kg (Kruskal-Wallis H test: *p*=0.003), 1.0 g/kg (Kruskal-Wallis H test: *p*=0.0002), and 2.0 g/kg (Kruskal-Wallis H test: *p*=0.0003) relative to age-matched vehicle-treated CONs. Glycyrrhizic acid did not affect intoxication ratings in CONs, but did restore the AIE-induced low intoxication rating to CON levels at ethanol doses of 0.5 g/kg (Kruskal-Wallis H test: *p*=0.012), 1.0 g/kg (Kruskal-Wallis H test: *p*=0.017), and 2.0 g/kg (Kruskal-Wallis H test: *p*=0.055) relative to age-matched vehicle-treated AIE animals. **(C)** Assessment of body temperature in adulthood across the cumulative ethanol challenge revealed increasing hypothermia with rising BECs regardless of treatment condition (main effect of Dose: *F*[3, 84]=331.5, *p*<0.0001). Prior AIE treatment in vehicle-treated rats decreased the hypothermic response across the cumulative ethanol challenge at doses of 0.5 g/kg (Bonferroni correction: *p*=0.018) and 1.0 g/kg (Bonferroni correction: *p*=0.003) relative to vehicle-treated CONs. Glycyrrhizic acid did not affect hypothermic responses in CONs, but it did reverse the AIE-induced blunting of hypothermia at ethanol doses of 0.5 g/kg (Bonferroni correction: *p*=0.015) and 1.0 g/kg (Bonferroni correction: *p*=0.017) relative to vehicle-treated AIE animals. **(D)** Assessment of balance on the tilting plane revealed a dose-dependent reduction in the angle of slide across treatment groups (main effect of Dose: *F*[3, 84]=336.9, *p*<0.0001). Prior AIE treatment in vehicle-treated rats increased the angle of slide at 0.5 g/kg (Bonferroni correction: *p*<0.0001), 1.0 g/kg (Bonferroni correction: *p*<0.0001), and 2.0 g/kg (Bonferroni correction: *p*<0.0001) relative to age-matched vehicle-treated CONs. While glycyrrhizic acid did not affect tilting plane performance in CONs, it restored cumulative ethanol-induced impairments at doses of 0.5 g/kg (Bonferroni correction: *p*=0.001), 1.0 g/kg (Bonferroni correction: *p*=0.009), and 2.0 g/kg (Bonferroni correction: *p*=0.005) relative to vehicle-treated AIE animals. **(E)** Assessment of loss of righting reflex (LORR) following the final dose of ethanol (i.e., 3.0 g/kg) revealed no differences across treatment conditions. **(F)** Assessment of plasma HMGB1 levels at baseline revealed that AIE treatment increased HMGB1 plasma levels relative to CONs (main effect of AIE: *F*[1, 26]=11.2, *p*=0.003). Follow-up analyses revealed significantly increased plasma HMGB1 levels in vehicle-treated AIE rats relative to vehicle-treated CONs (Šidák’s MCT: *p*=0.007) while glycyrrhizic acid treatment in AIE-treated animals blunted the increase in HMGB1 plasma levels. n=8 subjects/condition. Data are presented as mean ±SEM or median with interquartile range. \**p*<0.05, \*\**p*<0.01, \*\*\**p*<0.001, \*\*\*\**p*<0.0001, CON/Veh vs. AIE/Veh; $*p*<0.05, $$*p*<0.01, AIE/Veh vs. AIE/Glyz acid.

Assessment of plasma HMGB1 levels revealed that AIE increased HMGB1 plasma levels relative to CONs (main effect of AIE: *p*=0.003; **Figure 6F**). While the interaction was not significant, follow-up analyses revealed significantly increased plasma HMGB1 levels in vehicle-treated AIE rats relative to vehicle-treated CONs (*p*=0.007, Šidák) while glycyrrhizic acid treatment in AIE-treated animals blunted the increase in HMGB1 plasma levels. Previous studies have shown that AIE increases HMGB1 in the brain (39, 51, 58). The ERB endpoints include motor function assessments, so we performed immunohistochemistry in the M1 primary motor cortex. Immunohistochemical assessment of HMGB1 revealed a significant 1.2-fold increase in vehicle-treated AIE animals (*p*=0.0003, Šidák) that was restored to CON levels by glycyrrhizic acid (*p*=0.0005, Šidák; **Figure 7A**). Similarly, expression levels of phosphorylated (activated) NF-κBp65, which is a transcription factor involved in activation of proinflammatory signaling, was increased 1.4-fold in vehicle-treated AIE animals (*p*<0.0001, Šidák); this increase was reversed by adult glycyrrhizic acid treatment (*p*<0.0001, Šidák; **Figure 7B**). These findings suggest that blockade of HMGB1 by glycyrrhizic acid reverses AIE-induced neuroinflammation and persistent alcohol tolerance in adulthood.

**Figure 7.**
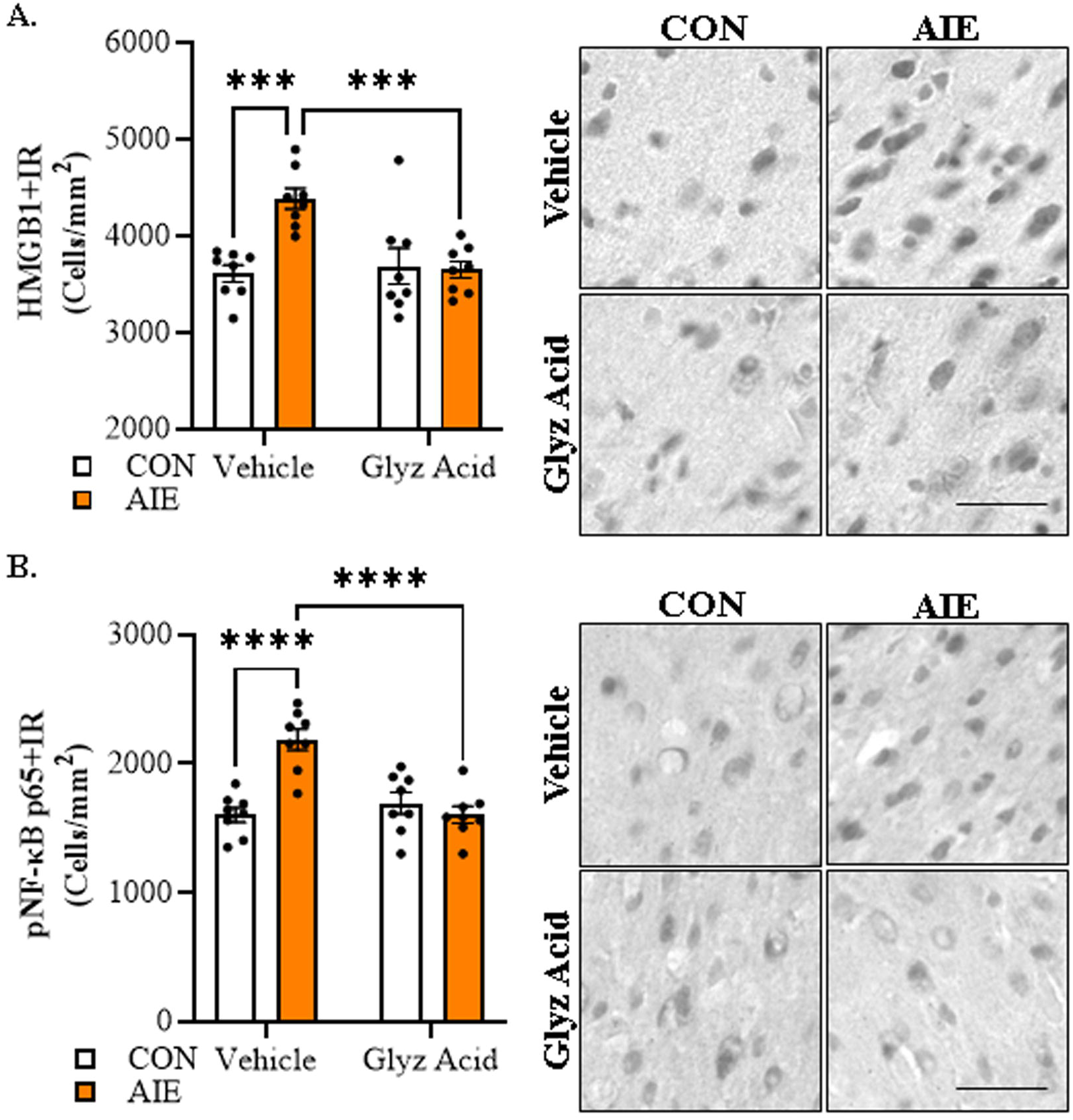
Immunohistological assessment of proinflammatory neuroimmune markers in the adult motor cortex following AIE treatment. **(A)** Immunohistochemical assessment of HMGB1+IR in the adult motor cortex revealed an approximate 1.2-fold increase in adult vehicle-treated AIE animals (Sidak’s MCT: *p*=0.0003), an effect that was reversed with glycyrrhizic acid (Sidak’s MCT: *p*=0.0005). **(B)** Immunohistochemical assessment of pNF-κBp65+IR in the adult motor cortex revealed an approximate 1.4-fold increase in adult vehicle-treated AIE animals (Sidak’s MCT: *p*<0.0001), an effect that was reversed with glycyrrhizic acid (Sidak’s MCT: *p*<0.0001). Scale bar = 30 μm. n=8 subjects/condition. Data are presented as mean ±SEM. \*\*\**p*<0.001, \*\*\*\**p*<0.0001.

## Discussion

To our knowledge, this is the first study to report that adolescent binge ethanol exposure confers lasting adult alcohol tolerance across a broad range of acute cumulative ethanol doses. The cumulative nature of the ERB allows dose-response assessments of alcohol responsivity as BECs progressively increase. Indeed, we observed that BECs were remarkably similar between treatment groups and across experiments suggesting that alterations in ethanol responsivity are not attributable to differences in ethanol metabolism. Alcohol tolerance is universally endorsed as a key symptom of AUD (2) and refers to diminished acute alcohol responsivity with continued use of the same quantity of alcohol. Although epidemiological studies link adult alcohol abuse and AUD with an adolescent age of drinking onset (6–11), the contribution of adolescent binge drinking to development of acquired tolerance in adulthood is poorly understood, due in part to the ethical considerations related to legal drinking age (65) and continued alcohol use across aging. Previous studies reported decreased alcohol responsivity in adult male rats following adolescent ethanol exposure on individual measures of motor function (23, 66, 67). We replicate and extend these studies to adult female Wistar rats and report that AIE reduces cumulative ethanol dose response impairments on measures of intoxication, hypothermia, tilting plane balance and LORR despite similar BECs. Further, we find baseline plasma levels of HMGB1 are persistently elevated in adult AIE-treated animals whereas HMGB1 plasma levels are significantly blunted following AIE across cumulative ethanol doses, consistent with diminished alcohol responsivity. These data suggest that adolescent binge drinking confers lasting alcohol tolerance, perhaps through a HMGB1 neuroimmune mechanism that may underlie the increased risk of AUD development associated with an adolescent age of drinking onset.

Emerging studies have identified neuroimmune activation as contributing to increases in alcohol drinking (41–44) and alcohol-induced neuropathology (19, 68–70). LPS (endotoxin) increases circulating plasma and brain levels of HMGB1 (71–73) and increases ethanol self-administration in rodents (74). Studies in AIE-treated adult rats as well as in the post-mortem human brain of individuals with AUD and an adolescent age of drinking onset find increased expression of neuroimmune signaling molecules, including HMGB1, throughout the brain (29, 39, 59, 69, 75–80). HMGB1 binds to and activates TLR4, RAGE, and other receptors, leading to nuclear translocation of NF-kBp65, thereby contributing to complex proinflammatory signaling (42, 62). Acute binge drinking and chronic alcohol abuse increase intestinal permeability, allowing gut bacterial endotoxin components (e.g., LPS) to enter systemic circulation, inducing proinflammatory signals that reach the CNS (60, 81), thereby altering cognition and drinking behaviors (82, 83). Our discovery that adolescent LPS mimics AIE-induced low ethanol responsivity and decreases plasma HMGB1 levels across cumulative ethanol doses in adulthood is consistent with inflammation contributing to the development of alcohol tolerance.

Glycyrrhizic acid, a constituent of *Glycyrrhiza glabra* (licorice) classified by the FDA as Generally Recognized As Safe as a food additive, is a selective HMGB1 inhibitor that binds to a pocket on HMGB1 blocking its proinflammatory activities through inactivation of released HMGB1 (64). We report post-AIE administration of glycyrrhizic acid restores adult alcohol responsivity to CON levels, blunts the persistent baseline induction of plasma HMGB1, and reverses HMGB1-mediated neuroinflammation in the primary motor cortex despite near-identical BECs compared to CONs at the time of testing. Our laboratory and others previously reported that anti-inflammatory therapeutics and exercise reverse AIE-induced neuropathology in adulthood (51, 84). Emerging studies suggest that AIE and HMGB1-mediated neuroinflammation can elicit chromatin remodeling in brain through epigenetic modifications, leading to changes in gene expression that contribute to alcohol-induced neuropathology that are reversible (29, 49, 52, 85–87). Reductions in acute responsivity across the cumulative ethanol challenge for different measures are likely dependent upon brain region, but may involve epigenetic alterations in neurocircuitry that are reversible. For instance, inhibition of histone deacetylases (HDACs) reverse development of rapid tolerance to the anxiolytic effects of alcohol (88). Further, alcohol drinking in alcohol-preferring rats has been linked to increases in HDAC activity (89). While the brain regional contributions to AIE-induced ethanol tolerance are likely related to individual ERB measures and remain to be fully elucidated, a recent study in adult mice found ethanol-induced rotarod impairments related to cerebellar ethanol metabolism-induced elevations of GABA levels (90). Our data suggest that AIE-induced developmental acquisition of alcohol tolerance in adulthood is reversible through blockade of HMGB1, providing a potential novel target for the development of therapeutics

In summary, we report AIE confers long-lasting alcohol tolerance in adulthood across a range of cumulative ethanol doses as well as blunted ethanol-induced HMGB1 plasma levels, consistent with neuroimmune tolerance. We report that adolescent LPS treatment, which models AIE-induced HMGB1-mediated neuroinflammation, confers lasting adult alcohol tolerance and blunted HMGB1 release across cumulative ethanol doses on the ERB, directly associating neuroinflammation with developmental acquisition of alcohol tolerance. Administration of the selective HMGB1 inhibitor glycyrrhizic acid reversed the AIE-induced low ethanol responsivity in adult AIE-treated animals, directly linking proinflammatory HMGB1 signaling to adolescent binge ethanol. These findings suggest a potential mechanistic target for the development of therapeutics for the treatment of AUD.

## Acknowledgements

Research reported in this publication was supported, in part, by the National Institute on Alcohol Abuse and Alcoholism (AA020024 (RPV/FTC), AA020023 (FTC), and AA019767 (FTC)) of the National Institutes of Health and the Bowles Center for Alcohol Studies. The content is solely the responsibility of the authors and does not necessarily represent the official views of the National Institutes of Health. **Fulton T. Crews:** Conceptualization, Methodology, Validation, Resources, Writing – Review & Editing, Visualization, Supervision, Funding Acquisition. All data generated in this study is available upon request. **Ryan P. Vetreno:** Conceptualization, Methodology, Validation, Formal Analysis, Investigation, Resources, Writing - Original Draft, Writing – Review & Editing, Visualization, Supervision, Funding Acquisition. We thank Jennie Vaughn for assistance editing the manuscript.

