## Supplement for "Adolescent Binge Ethanol Exposure Confers Lasting Alcohol Tolerance across a Cumulative Ethanol Challenge in Adulthood: Involvement of Proinflammatory HMGB1 Signaling"

***Supplemental Information***

### Supplemental Methods and Materials

### Ethanol Response Battery

#### *Behavioral Intoxication Rating Scale*

The ethanol intoxication rating scale has been used in binge drinking models for many years to provide an index of intoxication (1, 2). The 6-point behavioral intoxication rating scale was conducted as previously described (2, 3). Briefly, animals are scored by two researchers according to the following behavioral scale: (1) no sign of intoxication; (2) hypoactivity; (3) slight intoxication (ataxia; slight motor impairment); (4) moderate intoxication (obvious motor impairment; dragging abdomen); (5) high intoxication (dragging abdomen; LORR); (6) extreme intoxication (LORR; loss of eye blink response). The behavioral intoxication rating scale was conducted at baseline as well as 15 min after each ethanol dose for a total of five assessments.

#### *Body Temperature*

Hypothermia is a commonly studied ethanol response endpoint (4). Body temperature was assessed using a Thermalert clinical monitoring thermometer (Physitemp, Clifton, NJ) with an electric thermometer probe inserted approximately 5 mm into the rectum and left in place for ≥45 s until a stable reading was obtained. Animals were briefly restrained for 2 min in a DecapiCone (ThermoFisher Scientific, Austin, TX), and body temperature was assessed at baseline and again following each ethanol dose following completion of the behavioral intoxication rating scale for a total of five assessments. Difference (Δ) in body temperature as a consequence of cumulative ethanol dosing was calculated by subtracting body temperature following each ethanol dose from baseline body temperature. Room temperature was monitored daily and averaged 20.5°C (range 20°C to 21°C).

#### *Tail Blood Collection*

Tail blood was collected at baseline and 15 min after each administration of ethanol following body temperature assessment to determine BECs using a GM7 Analyzer (Analox; London, United Kingdom).

#### *Accelerating Rotarod*

The accelerating rotarod provides a measure of balance across cumulative doses of ethanol. Rats were trained on the accelerating rotarod (IITC Life Sciences, Woodland Hills, CA) for two days (three trials per day) prior to ERB assessment. The rotarod cylinder was 9.5 cm in diameter and 15 cm wide. On Training Day 1, each animal received three 3-min training trials with Trial 1 at 5 rotations per minute (rpm), Trial 2 at 10 rpm, and Trial 3 consisting of a start speed of 5 rpm and accelerating to 20 rpm over 3 min. If the animal fell off the rotarod cylinder during Training Day 1, the animal was gently placed back onto the cylinder for the remainder of training. Twenty-four h later on Training Day 2, each animal then received three 3-min training trials with Trial 1 at 10 rpm followed by two consecutive trials with the rotarod cylinder with a start speed of 5 rpm and accelerating to 20 rpm over 3 min. By the conclusion of rotarod training, nearly all female rats were able to remain on the rotarod cylinder for the entirety of the final 3-min training session.

Twenty-four h later, rotarod performance was assessed during the ERB, and each session consisted of three successive trials on the accelerating rotarod with a start speed of 5 rpm and accelerating to 20 rpm over 3 min. Latency to fall was recorded for every trial and was the primary outcome measure of the rotarod. Time (s) spent on the rotarod was assessed at baseline and again following each ethanol dose after completion of tail blood collection for a total of five sessions.

Each of the three successive trials per dosing session were averaged and change from baseline (Δ) in time spent on the rotarod as a consequence of cumulative ethanol dosing was calculated by subtracting time on the rotarod during baseline performance from each ethanol dose rotarod performance.

***Tilting Plane***

The tilting plane assesses motor coordination, which provides a measure of the ability of rodents to maintain balance as the angle of the horizontal plane is gradually increased. The tilting plane apparatus consisted of a clear Plexiglas box (60 cm × 24 cm × 20 cm) attached with a hinge to a frame with a glass panel floor. The box was tilted via an additional hinge attached to the base and a sliding protractor was used to measure the angle at which subjects began to slide down the glass floor panel. At the time of testing, the rat was placed on the apparatus facing away from the tilting hinge and the panel lifted slowly until the subject began to slide down the floor of the apparatus. The angle at which the rat began to slide was measured and the procedure repeated for three consecutive trials per session. The three trials within each session were averaged and difference (Δ) in angle of slide as a consequence of cumulative ethanol dosing was calculated by subtracting angle of slide from the averaged baseline angle of slide from each ethanol dose angle of slide.

***Loss of Righting Reflex***

Loss of righting reflex provides a measure of the sedative actions of ethanol as determined by the ability of a rat placed on its back to turn and upright itself. Loss of righting reflex was assessed following the final dose of ethanol after completion of the tilting plane. Animals were placed on their back in a V-shaped trough and assessed for righting reflex. Loss of righting reflex was defined as the inability of the rat to right itself onto all four paws within 60 s.

**Immunohistochemistry, Microscopic Quantification, and Image Analysis**

Animals were anesthetized with sodium pentobarbital (100 mg/kg, i.p.), transcardially perfused with 0.1 M PBS followed by 4.0% paraformaldehyde. Brains were excised and post-fixed in 4.0% paraformaldehyde for 24 hr at 4ºC followed by a 4 d fixation in 30% sucrose solution. Coronal sections were cut (40 µm) on a sliding microtome (MICROM HM450; ThermoScientific, Austin, TX, USA), and sections sequentially collected into well plates and stored at −20°C in a cryoprotectant solution (30% glycol/30% ethylene glycol in PBS).

Free-floating tissue samples containing the motor cortex (every $6^{th}$ section; approximate Bregma: 1.70 mm to -0.12 mm based on the atlas of Paxinos and Watson (5)) were washed in 0.1 M PBS, incubated in 0.6% $H_2O_2$ to inhibit endogenous peroxidases, and blocked with normal serum (MP Biomedicals). Sections were then incubated for 48 hr at 4ºC in a primary antibody solution containing blocking solution with either rabbit anti-HMGB1 (1:1000; Abcam, Cambridge, MA, USA, Cat. #ab18256) or rabbit anti-pNF-κB p65 (1:2000; Abcam, Cat. #ab86299). Sections were washed in PBS, incubated for 1 h in a biotinylated secondary antibody (Vector Laboratories), and incubated for 1 h in an avidin-biotin complex solution (1:200; Vector Laboratories). The chromogen nickel-enhanced diaminobenzidine (Sigma-Aldrich) was used to visualize immunoreactivity. Tissue was mounted onto slides, dehydrated, and cover-slipped. Negative controls for non-specific binding were conducted on separate sections employing the above-mentioned procedures omitting the primary antibody.

BioQuant Nova Advanced Image Analysis software (R&M Biometric, Nashville, TN) was used for image capture and quantification of immunohistochemistry. Representative images were captured using an Olympus BX50 microscope and Sony DXC390 video camera linked to a computer. For each measure, the microscope, camera, and software were background corrected and normalized to preset light levels to ensure fidelity of data acquisition. A modified unbiased stereological quantification approach was used, which was performed by experimenters blind to treatment, to quantify HMGB1+IR and pNF-κB p65+IR cells in the motor cortex. The corpus callosum was used as a landmark to assist with identification of the motor cortex. We previously reported that comparison of traditional unbiased stereological methodology with our modified unbiased stereological approach yielded nearly identical values for heterogeneously distributed cell populations (6). The outlined regions of interest were determined and data expressed as cells/mm$^2$.

**References.**

1. Obernier JA, White AM, Swartzwelder HS, Crews FT (2002): Cognitive deficits and CNS damage after a 4-day binge ethanol exposure in rats. *Pharmacology, biochemistry, and behavior*. 72:521-532.
2. Nixon K, Crews FT (2002): Binge ethanol exposure decreases neurogenesis in adult rat hippocampus. *Journal of neurochemistry*. 83:1087-1093.
3. Vetreno RP, Campbell J, Crews FT (2023): A multicomponent ethanol response battery across a cumulative dose ethanol challenge reveals diminished adolescent rat ethanol responsivity relative to adults. *Advances in Drug and Alcohol Research*. 3.
4. Silveri MM, Spear LP (2000): Ontogeny of ethanol elimination and ethanol-induced hypothermia. *Alcohol*. 20:45-53.
5. Paxinos G, Watson C (2014): *The rat brain in stereotaxic coordinates*. San Diego, CA: Academic Press.
6. Crews FT, Nixon K, Wilkie ME (2004): Exercise reverses ethanol inhibition of neural stem cell proliferation. *Alcohol*. 33:63-71.

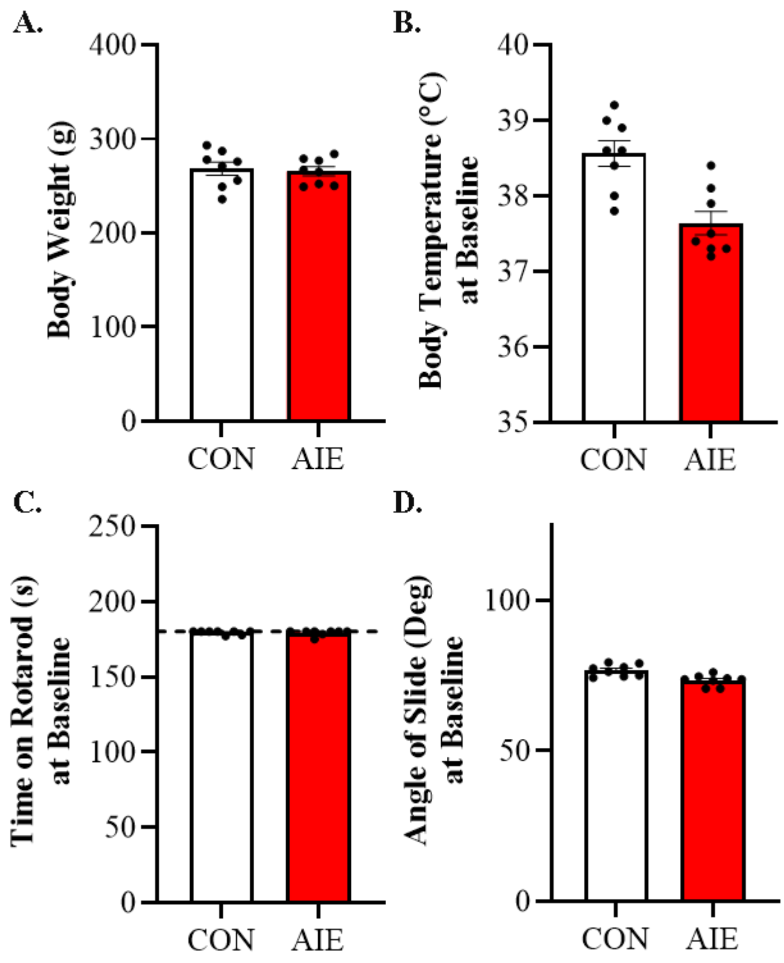


**Figure S1. Baseline measures during ethanol response battery (ERB) assessment for Experiment 1. (A)** Baseline body weights of CON- (268 g ±7.0) and AIE-treated (265 g ±5.0) animals at the time of ERB testing (i.e., P75). **(B)** Baseline body temperatures in CON- (38.5°C ±0.17) and AIE-treated animals (37.6°C ±0.16) at the time of ERB testing. **(C)** Baseline time (s) on the accelerating rotarod between CON- (179.3 s ±0.44) and AIE-treated (179.2 s ±0.63) animals. Dashed line indicates 3 min trial duration. **(D)** Baseline angle of slide on the tilting plant between CON- (76.7° ±0.68) and AIE-treated (73.4° ±0.65) animals. n=8 subjects/condition. Data are presented as mean ±SEM.

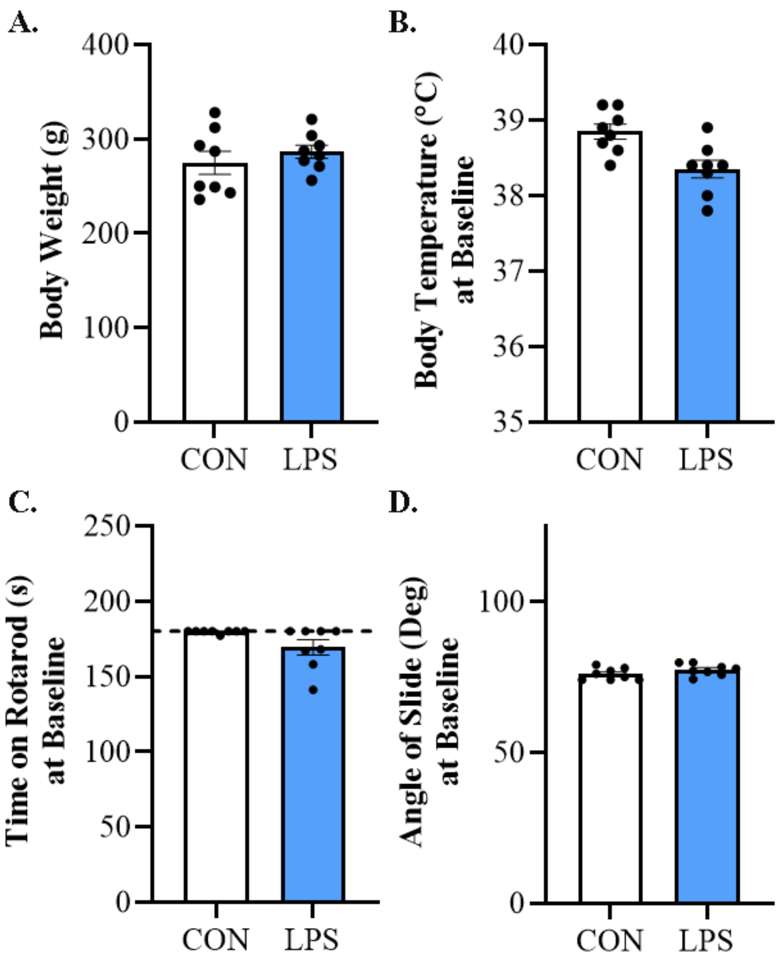


**Figure S2. Baseline measures during ethanol response battery (ERB) assessment for Experiment 2. (A)** Baseline body weights of CON- (275 g ±12.3) and LPS-treated (287 g ±7.0) animals at the time of ERB testing (i.e., P80). **(B)** Baseline body temperatures in CON- (38.9°C ±0.10) and LPS-treated animals (38.4°C ±0.12) at the time of ERB testing. **(C)** Baseline time (s) on the accelerating rotarod between CON- (179.6 s ±0.38) and LPS-treated (169.3 s ±5.0) animals. Dashed line indicates 3 min trial duration. **(D)** Baseline angle of slide on the tilting plant between CON- (75.9° ±0.69) and AIE-treated (77.4° ±0.66) animals. n=8 subjects/condition. Data are presented as mean ±SEM.

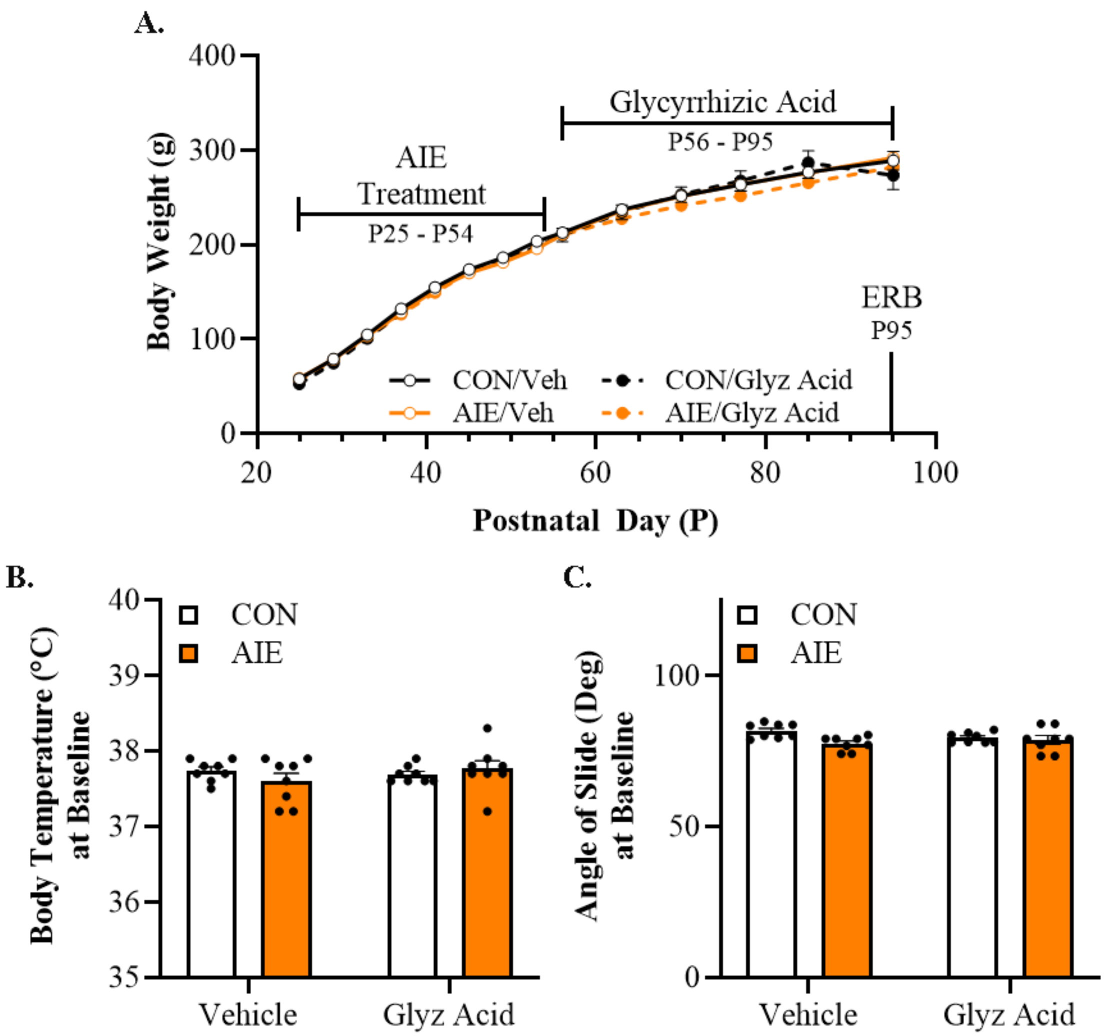


**Figure S3. Baseline measures during ethanol response battery (ERB) assessment for Experiment 3. (A)** While female Wistar rats evidenced dramatic body weight gains across Experiment 3, neither AIE or glycyrrhizic acid (glyz acid) treatment affected body weights. **(B)** Baseline body temperatures in CON- (Vehicle: 37.7°C; Glyz Acid: 37.7°C) and AIE-treated animals (Vehicle: 37.6°C; Glyz Acid: 37.7°C) at the time of ERB testing (i.e., P9). **(C)** Baseline angle of slide on the tilting plant between CON- (Vehicle: 81.7°; Glyz Acid: 79.3°) and AIE-treated (Vehicle: 77.4°; Glyz Acid: 78.6°) animals. n=8 subjects/condition. Data are presented as mean ±SEM.
